# DNA-binding domain-aware classification enables systematic annotation of the regulatory genome

**DOI:** 10.1101/2024.12.20.629625

**Authors:** Christian Ickes, Cigdem Hazal Timucin, Bendix Christian Harms, Umut Akgül, Íñigo Vicente-Hernández, Ivan Bogeski, Tim Beißbarth, Martin Haubrock

## Abstract

Defining the cis-regulatory code remains one of the central challenges of modern genomics, requiring the reliable association of transcription factor binding sites (TFBSs) with their cognate transcription factors (TFs) from DNA sequence information. While deep learning methods outperform conventional position weight matrix (PWM)-based approaches, both typically formulate TFBS prediction as a bound-versus-unbound decision rather than resolving the competitive binding potential among TFs at a given genomic location. This is further complicated by the widespread sharing of DNA-binding domains (DBDs) among TFs, which produces overlapping binding preferences, rendering TFBS assignment at the individual TF level inherently ambiguous.

Consequently, we present TFClassPredict, a DNABERT-based framework that reframes TFBS prediction as a multi-class classification problem across 23 DBD-classes, discriminating each against all remaining DBD-classes, directly enabling the resolution of binding events between DBD-classes. Trained on high-confidence directly bound TFBSs and exploiting DBD-DNA co-evolution, TFClassPredict naturally resolves the ambiguity arising from shared DBD architectures.

We demonstrate that TFClassPredict achieves robust DBD-class level classification, outperforming PWM-based approaches and benchmarked deep learning architectures with predictions mapping to biologically defined DBD-classes. Beyond classification, TFClassPredict improves PWM-based pipelines, three-dimensional genome interaction prediction, and allows genome-wide DBD-class annotation, while recapitulating known lineage-specifying TF programs from immune cell ATAC-seq data.

## 1 Introduction

Transcription factors (TFs) are DNA-binding proteins that play central roles in the regulation of gene expression [1]. Understanding TF–DNA recognition is essential for dissecting cellular processes, cell-fate decisions, and cell development [2– 4], but also disease-associated regulatory dysregulation [5]. The short DNA sequences bound by TFs are referred to as transcription factor binding sites (TFBSs), which often co-occur to form cis-regulatory modules (CRMs) [6]. The composition and spatial arrangement of TFBSs within CRMs are critical determinants of gene expression dynamics [7]. Therefore, reliably predicting potential TFBSs is essential for interpreting the regulatory genome.

Computational TFBS prediction conventionally relies on position weight matrices (PWMs) [8]. Although widely used and interpretable, PWMs represent TF binding preferences as independent contributions of individual nucleotide positions within a short fixed-length motif, limiting their ability to model higher-order sequence dependencies, variable motif architectures, and contextual effects from flanking nucleotides [9, 10]. Deep learning approaches, including Convolutional Neural Networks (CNNs) and Recurrent Neural Networks (RNNs), have improved TFBS predictions by learning nonlinear and context-dependent sequence patterns directly from data [11–14]. More recently, transformer-based genomic language models such as DNABERT have further extended this paradigm by representing DNA sequences as overlapping k-mers and learning contextual sequence representations through selfattention, achieving improved performance on TFBS prediction tasks [15, 16]. However, these models typically frame TFBS prediction as a binary bound-or-unbound decision for a single TF, leaving unaddressed the biologically more relevant question of which TF holds the dominant binding potential at a given genomic location. This distinction is of particular relevance for understanding the compositional logic of CRMs, where the identity and relative binding potential of co-occurring TFs ultimately determine regulatory specificity and outcome.

TF–DNA binding is commonly mediated by two complementary mechanisms: base read-out and shape readout [17, 18]. Base read-out relies on recognition of specific nucleotide sequences, whereas shape readout captures sequence-dependent three-dimensional DNA features. DNA-binding domains (DBDs) adopt diverse structural architectures [19], including helix–turn–helix, zinc finger, and basic leucine zipper domains, among others, and constitute major determinants of DNA-binding specificity [20, 21]. These structural relationships have been systematically cataloged in TFClass [22], a hierarchical classification system that organizes TFs according to their DBD architecture. DBD structure and DNA binding site sequence preference are mutually constrained through co-evolution, such that TFs sharing similar DBD architectures consistently recognize similar binding site sequences [23–25]. Studies have shown that incorporating DBD structural similarity among TFs improves TFBS prediction [26–28], while resolving TF binding below the DBD level remains particularly challenging [26].

This challenge motivates the DBD-class level as a biologically valid abstraction for analyzing transcription factor binding potential (TFBP), assessing the capacity of each DBD-class to specifically and directly engage regulatory regions as a natural unit of sequence-based regulatory annotation (Figure 1a). We investigate whether this DBD-class level abstraction is learnable from DNA sequence alone, contrasting the inherent limitations of position-independent PWM-based approaches with a deep learning strategy based on DNABERT, a k-mer-based transformer pre-trained on genomic sequences. Thereby, we develop TFClassPredict (Figure 1b), a framework for inferring TFBP stratified by DBD-class, finetuned on high-confidence TFBSs derived from UniBind [29], a comprehensive resource of direct TF-DNA interactions generated by uniformly processing thousands of ChIP-seq datasets. We demonstrate that TFClassPredict accurately classifies TFBSs across 23 DBD-defined TF classes, complementing PWM-based tools for refined TF binding site identification and enabling more complex downstream regulatory analyses. Furthermore, we show that TFClassPredict enables genome-wide characterization of DBD-class level TF binding preferences and show its utility in a biological application by recovering known TF regulatory programs underlying immune cell differentiation from ATAC-seq data. To facilitate broad community adoption, TFClassPredict is provided as a user-friendly classification application together with a genome-wide DBD-class binding potential annotation tracks for the human genome (Figure 1c).

**Fig. 1.**
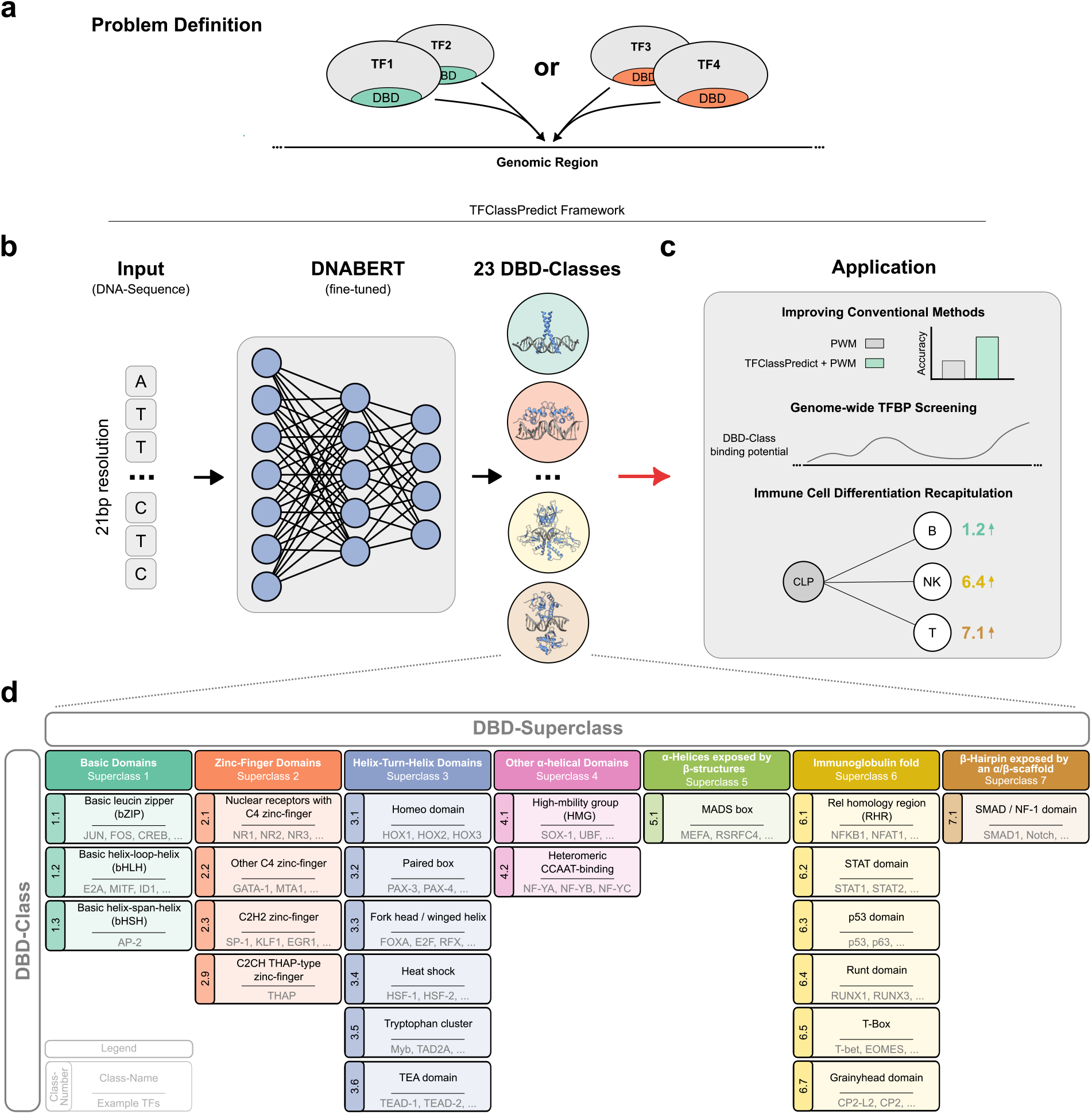
The TFClassPredict framework for reliable TFBS-to-DBD association. **a:** Conventional TFBS prediction methods frame binding as a binary bound-or-unbound decision for a single TF, and the widespread sharing of DBD architectures among TFs further complicates the unambiguous assignment of TFBSs, leaving unaddressed which TF holds the dominant binding potential at a given genomic location, a distinction critical for understanding the compositional logic of CRMs. **b:** TFClassPredict reframes TFBS prediction as a multi-class classification problem across 23 DBD-classes, directly resolving binding ambiguity arising from shared DBD architectures and enabling discrimination between TF groups. **c:** TFClassPredict is applicable across a range of downstream analyses, including the improvement of PWM-based pipelines, genome-wide DBD-class annotation, three-dimensional genome interaction prediction, and the recapitulation of known lineage-specifying TF programs from immune cell ATAC-seq data. **d:** TFClassPredict employs 23 TFClass-derived DBD categories to predict TFBPs, grouped into seven higher-level superclasses, with representative TFs shown for each category.

## 2 Results

### Conventional approaches fail to reliably recover biologically validated TFBSs among DBD-classes

Inferring the regulatory landscape and regulatory differences based on sequencing strategies for open chromatin regions relies heavily on recovering TF-DNA interactions from sequencing data. This is conventionally achieved by PWM-based approaches for individual TFs, which further depend on appropriate negative sets representing sequences unlikely to be bound by TFs, typically generated by nucleotide shuffling or GC-matched genomic background sampling [30, 31]. DBD similarity has been shown to directly predict DNA-binding specificity [23], and the highest-scoring PWM for a ChIP-seq peak is typically derived from the same DBD-class [26], suggesting that PWM-based approaches should in principle be capable of reliably identifying directly bound TFBSs at the DBD-class level. We therefore systematically assessed this capacity across 23 DBD-classes using high-confidence directly bound TFBSs from UniBind (Figure 1d).

For a robust evaluation of models discovering TF-DNA interactions, a reliable data source is necessary. To that end, we leverage TFBSs annotated in the UniBind database [29], which processes thousands of ChIP-seq datasets with the ChIP-eat pipeline to obtain millions of high-confidence, directly bound TFBSs (Supplement Figure A1a) across several DBD-classes. We expanded all binding sites to a full 21bp window, corresponding to the largest binding site in the dataset (Supplement Figure A1b). To ensure that only single binding sites are present within each window, we further removed TFBSs that showed overlapping binding sites (Supplement Figure A1c).

We applied HOMER [32], a de novo motif enrichment analysis tool, to identify motifs within the stack of TFBSs associated with individual DBD-classes. While HOMER (Figure 2a) consistently identified PWMs corresponding to the queried DBD-class, it suffered from a high number of false-positives, with the majority of enriched motifs incorrectly assigned to the queried DBD-class and only a minority corresponding to the expected DBD-class level. Similarly, FIMO [33], a widely used PWM-based motif scanning algorithm, applied to individual sequences using JASPAR-derived PWMs identical to those used in UniBind processing, yielded consistently and substantially low classification accuracies across all 23 DBD-classes (Figure 2b). Collectively, these results demonstrate the inherent limitations of PWM-based approaches for TFBS classification, even when evaluated at the DBD level.

**Fig. 2.**
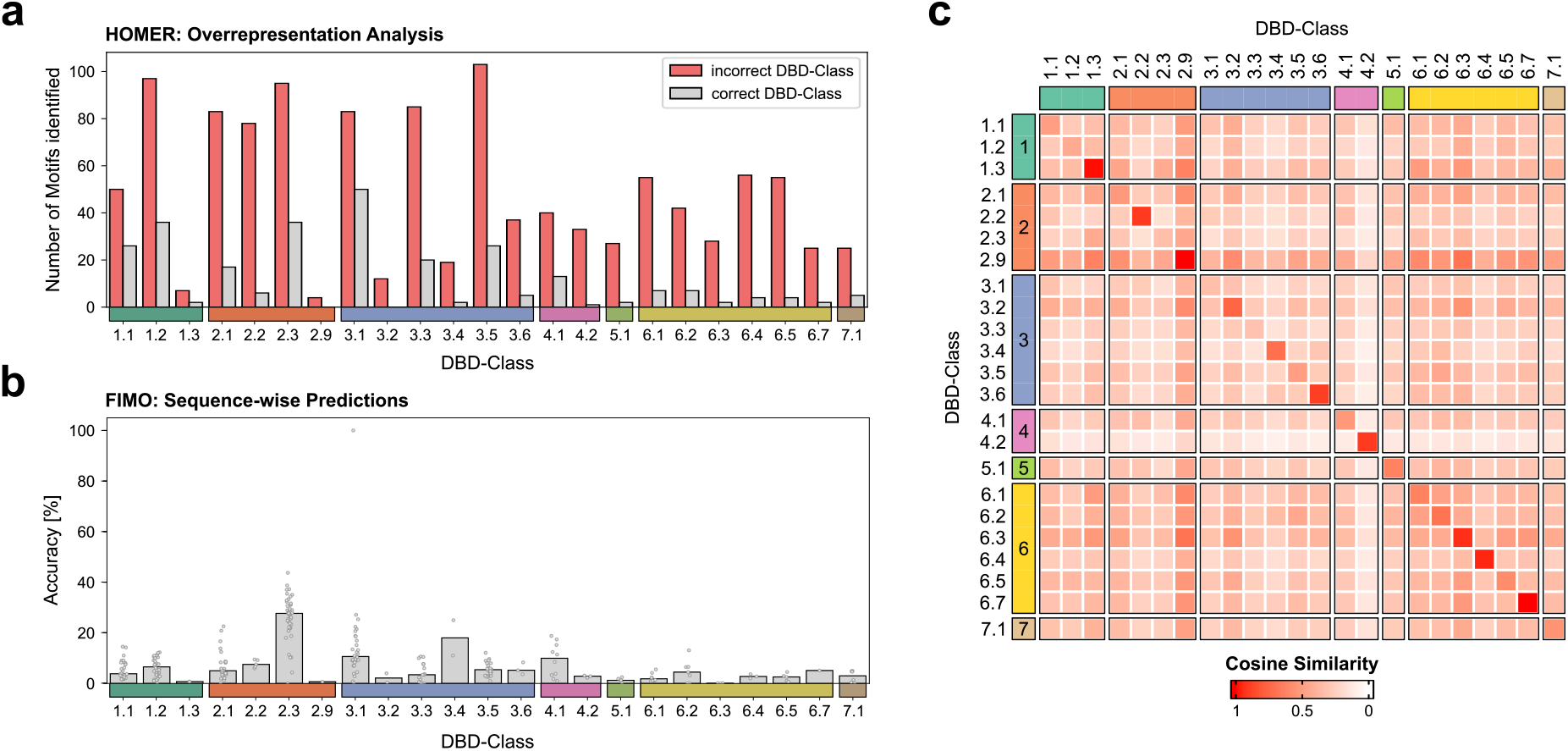
PWM-based sequence analysis demonstrates diminishing performance even across highly similar DBD-classes. **a:** HOMER overenrichment analysis based on non-overlapping TFBSs per DBD-class on the UniBind test set. Identified motifs demonstrate high false-positive rates across all DBD-classes. **b:** Sequence-wise predictions on the UniBind test set were performed with FIMO. Dots represent TF-wise accuracies; bars indicate the median accuracy across TFs. **c:** Cosine similarity within the 6-mer space between non-overlapping TFBSs per DBD-class, averaged across TF-wise similarities. Row- and column-wise color codes indicate the DBD-superclass level.

To validate the biological coherence of the DBD-class sequence grammar in our dataset, we computed DBD-class-wise cosine similarities of 6-mer frequencies across TFBSs by averaging TF-wise cosine similarities within each DBD-class. The resulting heatmap reveals the highest similarity within each DBD-class, reflecting the conserved DNA recognition of related DBD architectures. A notable degree of cross-class similarity is observable across all classes, consistent with the well-documented convergence of core recognition sequences across structurally distinct DBD-classes [5, 34].

Taken these findings together, the positional independence assumption, the reliance on reliable background sets, and the single-consensus representation inherent in PWM-based approaches collectively impose an upper bound on classification performance at the DBD-class level. The cross-class k-mer similarity, and by extension the cross-TF similarity demonstrated in Figure 2c, constitutes an irremediable source of misclassification [8, 10], irrespective of the specific tool or analytical strategy employed, and is consistent with the high false positive rates associated with conventionally PWM-based binding site scanning. The HOMER and FIMO analyses provide converging evidence that PWM-based motif enrichment at default parameters lacks the discriminative power to reliably distinguish binding sites, with elevated false positive rates posing a particular challenge for tools that rely solely on sequence information to analyze regulatory dynamics.

### TFClassPredict employs DNABERT to deliver robust DBD-class level binding site classification

As demonstrated in the preceding analyses, current approaches for recovering TFBSs from sequence information alone have substantial limitations. While restricting predictions to the DBD level already improves performance, these gains remain insufficient to reliably distinguish binding sites. Overcoming these limitations requires a fundamentally different approach that captures the full complexity of DBD-class level sequence grammar beyond the constraints of position-independent PWM models. Deep learning models represent a promising strategy in this regard, offering the capacity to learn context-dependent sequence features that are inaccessible to conventional PWM-based methods.

To identify the optimal model architecture for classifying TFBSs by DBD-class, we systematically evaluated multiple deep learning architectures using the UniBind repository of high-confidence, non-overlapping TFBSs spanning 23 DBD-classes. We compared two established models, namely DNABERT, a k-mer-based transformer, and DanQ, an RNN operating on one-hot encoded inputs, against three baseline models: a Neural Network (NN) operating on k-mer-tokenized sequences, a NN operating on one-hot encoded sequences, and a weighted random baseline informed by class-frequency priors. The classification task was formulated as a multi-class problem, in which each DBD-class is distinguished from all remaining classes simultaneously, thereby circumventing the background sampling biases inherent to binary classification approaches. Model performance was assessed using accuracy and macro-averaged F1 score across all DBD-classes, complemented by per-TF accuracy.

Performance evaluation revealed substantial differences across all tested architectures (Figure 3a). The weighted random baseline, with accuracies and F1 scores as low as ∼10%, confirmed the inherent difficulty of the classification task and established the minimum threshold any meaningful classifier must exceed. Notably, even simple NN architectures demonstrated that k-mer tokenization substantially outperforms one-hot encodings, highlighting the critical importance of sequence representation for TF binding classification. Among all evaluated architectures, DNABERT achieved the highest overall classification performance (Accuracy: 87,3%, F1_macro_: 78,1%; Supplementary Table A2), with context-aware sequence models (DNABERT, DanQ) broadly outperforming simpler baselines. The superiority of DNABERT can be attributed to its combination of k-mer-based tokenization, self-attention mechanism, and pretraining on the human genome, properties that are particularly well-suited to capturing the combinatorial and context-dependent sequence grammar underlying TF binding. Per-TF accuracy analysis further confirmed this superiority, with DNABERT achieving higher median accuracy than all alternative architectures across the majority of DBD-classes, out-performing DanQ in 19 of 23 classes (Figure 3b, Supplementary Table A2).

**Fig. 3.**
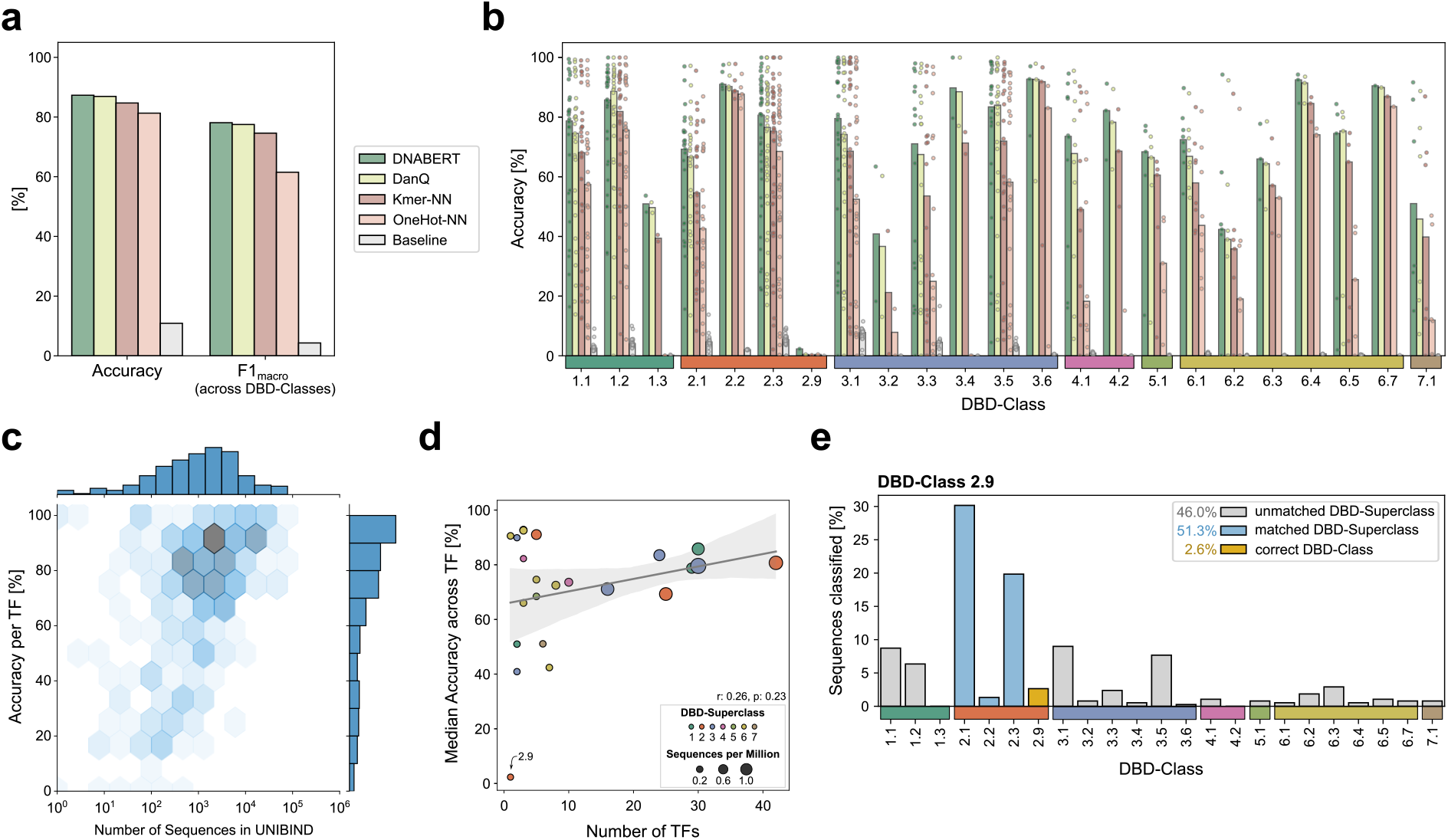
TFClassPredict model selection and performance evaluation using UniBind data identifies DNABERT as the best-performing model architecture. We evaluated multiple model architectures to assess classification performance across DBD-classes. **a:** Accuracy and F1 score averaged across DBD-classes for the different model architectures tested. **b:** Accuracy for individual TFs. Each dot represents a single TF, and bars indicate the median TF-wise accuracy for each class and model architecture. **c:** DNABERT-based accuracy for individual TFs compared to the number of sequences available per TF. **d:** DNABERT-based median accuracies across TFs per DBD-class **e:** Sequence classification results for DBD-class 2.9.

A key practical consideration for DBD-class level classification is the variability in the number of TFBSs available per TF across DBD-classes, reflecting the well-documented non-uniformity of ChIP-seq data coverage across the TF repertoire (Supplement Figure A1a). When examining DNABERT per-TF accuracy as a function of available training sequences, we observed, as expected, a positive relationship between the number of training sequences and classification accuracy (Figure 3c). Importantly, as demonstrated in Figure 3d, pooling binding site data across all TFs within a given DBD-class allows DNABERT to effectively mitigate the limitations imposed by low TF representation. This strategy maintains robust classification performance even for DBD-classes characterized by sparse TF representation, as evidenced by a weak and statistically insignificant association between TF number and classification accuracy (Pearson’s *r* = 0.26, *p*-value = 0.23, *N*_DBD-classes_ = 23). Nevertheless, DBD-classes with both low training sequence counts and low TF numbers remain challenging, as exemplified by DBD-class 2.9, which continues to exhibit diminished classification performance under these conditions.

Finally, to illustrate the sequence-level classification behavior of DNABERT, we present detailed classification results for the most challenging DBD-class 2.9 as a representative example (Figure 3e). DBD-class 2.9 represents a particularly difficult case, as it is represented by only a single TF with a limited number of binding sequences, and exhibits high k-mer similarity (Figure 2c) to other DBD-classes, making it inherently difficult to distinguish at the sequence level. Nevertheless, more than 50% of misclassified sequences are attributed to their corresponding DBD-superclass, a pattern consistently observed across all classes (Supplementary Figure A2a) and exceeding random expectation (Supplementary Figure A2b), demonstrating that the DNABERT-based classifier retains biologically meaningful classification behavior even in challenging cases, correctly resolving TFBSs at the DBD-superclass level across a diverse set of binding site sequences.

These results demonstrate that DNABERT-based DBD-class level aggregation effectively decouples classification performance from per-TF data availability, establishing DNABERT as the underlying model of TFClassPredict and a robust solution for DBD-class binding site classification across the full diversity of the human TF repertoire.

### TFClassPredict enhances prediction robustness across diverse analytical applications

Having characterized the classification performance of TFClassPredict, we next assessed its utility as a practical filtering step for existing PWM-based binding site prediction pipelines. Specifically, we evaluated whether applying TFClassPredict as a post-hoc filtering step to FIMO-based binding site predictions could reduce false positives and improve overall classification accuracy. Its application to FIMO predictions resulted in a substantial improvement in median per-TF classification accuracy across all DBD-classes (Figure 4a), confirming that TFClassPredict effectively identifies and removes false positive FIMO predictions that are inconsistent with the expected DBD-class level sequence grammar. A representative example for DBD-class 2.3 illustrates this filtering effect at the individual DBD-class level, demonstrating that the improvement in classification accuracy is consistent and reproducible across individual TFs belonging to the same DBD-class (Figure 4b).

**Fig. 4.**
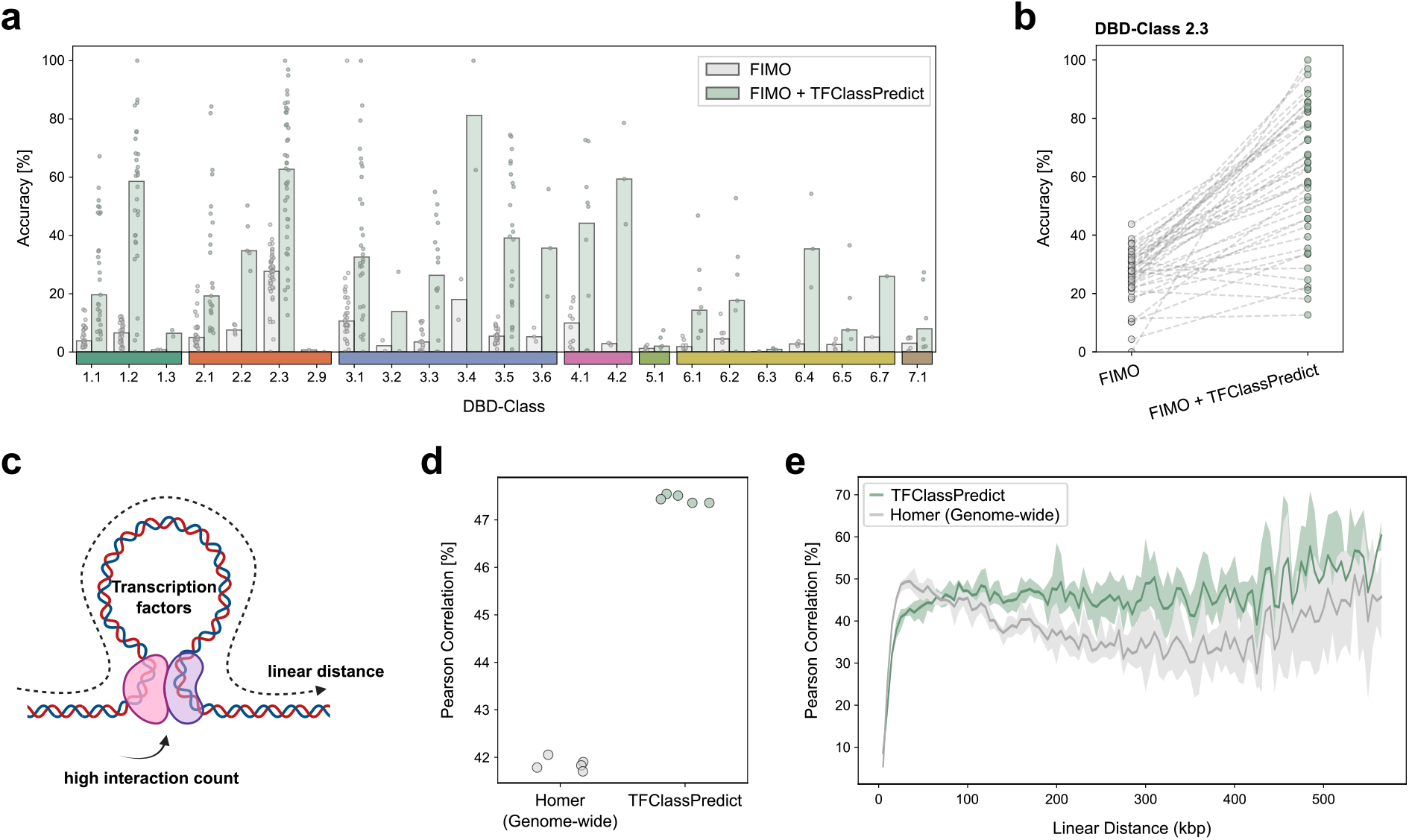
TFClassPredict enhances robustness across diverse analytical strategies. Applying TFClassPredict as a filtering step to FIMO predictions reduces false positives and improves prediction accuracy. **a:** Median accuracy across TFs before and after filtering FIMO predictions with TFClassPredict. **b:** Representative example for DBD-class 2.3. To further assess its utility in a more complex setting, we applied TFClassPredict to predict three-dimensional genome interactions from Micro-C of H1-hESC cells using TFBPs located within ATAC-seq peaks. **c:** Schematic illustration of the genome interaction framework. (Created with BioRender) **d:** Pearson correlation between observed and predicted distance-corrected interaction counts for RF models based on either HOMER-predicted or TFClassPredict-predicted TFBP. **e:** Pearson correlation coefficient stratified by linear genomic distance (bin size: 5kb) for RF models trained on HOMER-derived versus TFClassPredict-derived TFBPs. Shaded areas indicate the minimum and maximum of the Pearson correlation coefficient within the corresponding bin across the five cross-validation folds.

To further assess the utility of TFClassPredict in a more complex and biologically relevant analytical setting, we applied it to predict three-dimensional genome interactions from Micro-C data. To this end, we estimated transcription factor binding potentials (TFBPs) using TFClassPredict, where each TFBP reflects a potential binding site by TFs belonging to a given DBD-class. The number of TFBPs falling within defined genomic regions derived from ATAC-seq peaks serves as a quantitative measure of DBD-class level binding potential, integrating both sequence-based binding predictions and chromatin accessibility information. These TFBPs were used as features in Random Forest (RF) models to predict distance-corrected chromatin interaction counts (Figure 4c). We compared two RF models: the first used TFBPs derived from PWMs identified by HOMER, mapped to their corresponding TFs and assigned to DBD-classes via TFClass, and the second used TFBPs derived directly from TFClassPredict. Both models were otherwise methodologically matched, enabling a direct comparison of the two feature sets.

The RF based on TFClassPredict-derived TFBPs achieved increased Pearson correlation between observed and predicted distance-corrected interaction counts compared to the RF based on HOMER-derived binding potentials (Figure 4d), consistent over two different cell lines (Supplement Figure A3, Supplementary Table A3), demonstrating that the higher quality DBD-class level binding site predictions of TFClassPredict translate into enhanced performance on the downstream task of three-dimensional genome interaction prediction. Stratification by linear genomic distance revealed a nuanced distance-dependent performance pattern (Figure 4e). At short genomic distances (0-60 kbp), HOMER-derived binding potentials achieved higher Pearson correlation, possibly reflecting small, non-extruded loop domains, much smaller than a typical topologically associated domain (TAD), which are dominated by CTCF binding sites and other structural components of the cohesin complex, such as RAD21 and SMC3. These are commonly referred to as CTCF loop anchors and are well captured by the CTCF PWM [35, 36]. Beyond 60 kbps, however, TFClassPredict-derived DBD-class level binding potentials achieved significantly higher Pearson correlations, possibly reflecting the increasing dominance of long-range functional enhancer-promoter contacts [35]. The higher DBD binding potential activity observed at longer distances therefore might reflect a genuine shift from structurally dominated to TF-binding grammar-driven chromatin interactions, establishing TFClassPredict as particularly well-suited for predicting functional long-range regulatory contacts.

These results establish TFClassPredict not only as a standalone classifier but as a broadly applicable tool for enhancing the reliability of existing PWM-based binding site prediction workflows, as well as for denoising predictions in more complex downstream analyses.

### Genome-wide mapping of DBD-class binding potentials reveals class-specific enrichment across regulatory regions and functional associations

Having demonstrated the classification performance and analytical utility of TFClassPredict, we next applied it at genome scale to generate a comprehensive map of DBD-class specific TFBPs across the human genome. For each of the 23 DBD-classes, TFClassPredict was applied genome-wide, providing, to our knowledge, the first systematic annotation of the human genome by DBD-class binding site grammar.

The analysis revealed a non-uniform distribution of binding potentials across the DBD-classes (Figure 5a). The majority of TFBPs were assigned to DBD-class 2.1, corresponding to nuclear receptors with C4 zinc fingers, consistent with the known abundance and regulatory importance of this DBD-class in the human genome [5, 37]. Superclass level annotation of the outer ring confirmed that zinc-fingers (DBD-superclass 2) and helix-turn-helix domains (DBD-superclass 3) account for the largest proportion of predicted TFBPs across the human genome, reflecting the dominance of these DBD architectures in human gene regulation. Analysis of TF frequencies stratified by superclass annotation further confirmed this distribution, with DBD-superclasses 2 and 3 again representing the most frequently predicted classes across the human TF repertoire (Figure 5a, right panel).

**Fig. 5.**
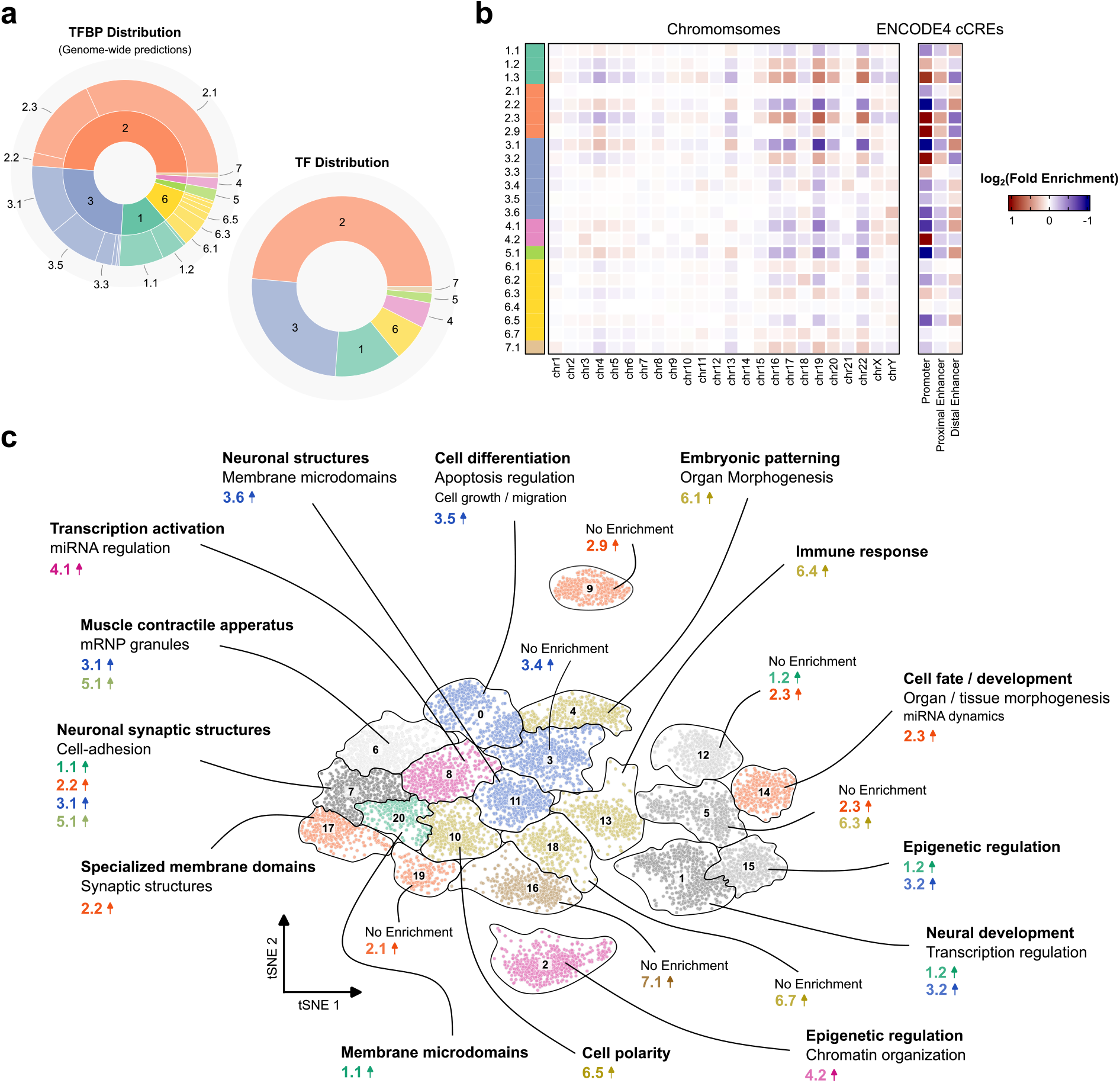
Genome-wide prediction of DBD-class binding potentials reveals DBD-class specific regulatory region enrichments. **a:** Distribution of TFBPs for individual DBD-classes estimated by TFClassPredict (left) and the number of TFs per DBD-superclass from Wingender et al. [22] (right). **b:** Fisher’s exact test enrichment analysis resolved by chromosome (left) and ENCODE4 cCREs (right), showing log_2_ fold enrichment relative to all other entries as background. **c:** t-SNE plot of ENCODE4 cCREs-defined promoter regions within the top 1% of transcription factor binding potential deviations (TFBPDs) in at least one DBD-class, clustered by agglomerative hierarchical clustering. Cluster annotations reflect enriched Gene Ontology (GO) terms depending on their corresponding sequence features. Arrows indicate the DBD-class in which the cluster were significantly enriched, exhibiting higher values in their TFBPDs compared to all other selected promoters, as determined by one-sided Mann-Whitney U tests (alternative: greater) with Bonferroni correction applied, retaining only features of high stringent significance (*p*_*adj*_ < 10^−100^).

To characterize the relationship between DBD-class specific binding potentials and defined genomic elements, we summarized them by chromosome (Figure 5b, left heatmap; Supplementary Table A4) or by ENCODE-defined regulatory regions comprising three histone modification-defined categories (promoters, proximal enhancers, and distal enhancers; Figure 5b, right heatmap; Supplementary Table A4). While no strong chromosome-related bias was observed, regulatory-relevant regions indicated much more distinct patterns. Strikingly, promoter regions showed the most pronounced and informative DBD-class specific enrichment. Notable examples of promoter-enriched DBD-classes include the strong promoter enrichment of NF-Y binding potentials (DBD-class 4.2), consistent with the well-established role of the CCAAT box as one of the most conserved promoter-proximal motifs in eukaryotic genes, typically located 60-100 bp upstream of the transcription start site (TSS) [38], and the pronounced promoter-proximal occupancy of paired-box TFs such as PAX5 (DBD-class 3.2), which regulates target gene expression through promoter-proximal recruitment of chromatin-modifying complexes [39]. In contrast, proximal and distal enhancers showed less pronounced differences in DBD-class binding potentials, suggesting that promoter regions are more deterministically constrained in their DBD-class composition than distal regulatory elements. This observation is in line with the known preference of certain DBD-classes for distal regulatory elements. GATA factors (DBD-class 2.2), MADS-domain TFs (DBD-class 5.1) such as MEF2A and SRF, and homeodomain TFs (DBD-class 3.1) covering HOX, all show a pronounced depletion of binding potentials at proximal promoters (Figure 5b), instead predominantly occupying distal regulatory, intronic, and intergenic regions. This distribution is consistent with their well-documented roles in enhancer-mediated gene regulation, ranging from hematopoietic and developmental control by GATA factors [40] to signal-responsive enhancer activation by MEF2A and SRF [41] to distal regulatory binding by HOX factors such as HOXA2 in developmental gene regulation [42]. The consistency of DBD-class enrichment profiles across chromosomes further confirms that the observed patterns reflect genuine constraints of regulatory sequence grammar rather than biases in chromosomal or regional sequence composition.

To analyze single genomic regions, we defined transcription factor binding potential deviations (TFBPDs) as a well-normalized scoring system to evaluate differences in binding potential while accounting for region length and DBD-class frequency, allowing between-class and between-region comparisons (see Methods 4.7). The definition of TFBPDs enabled exploration of the functional organization of DBD-class binding potentials in promoter regions. To that end, we embedded all human promoters retaining a top 1% TFBPD in at least one DBD-class (*N*_promoters_ = 9317) into a two-dimensional t-SNE space (Figure 5c) based on their TFBPD profiles, and selected the best clustering based on the optimal silhouette score (Supplementary Table A4). The resulting embedding revealed distinct clusters of promoters defined by their DBD-class binding potential composition, demonstrating that promoters can be systematically organized by their DBD-class level sequence grammar (Supplement Figure A4). Cluster annotation based on DBD-class over-enrichment revealed biologically coherent associations between DBD-class binding potentials and the functional identity of regulated genes (Supplementary Table A4). At the global level, promoters enriched for DBD-class binding potentials were predominantly associated with cell development processes and cell-fate determination. More specifically, DBD-class 3.6, corresponding to TEA-domain/TEF-1 factors, showed enrichment of binding potentials near genes associated with neuronal structures and membrane-related annotations. This finding aligns with the established role of TEAD factors as DNA-binding partners of YAP in vertebrate neural tube development, where YAP/TEAD signaling governs neural progenitor proliferation, differentiation, fate choice, and survival [43]. Similarly, DBD-class 6.4, corresponding to Runt-domain factors, showed enrichment of binding potentials near genes associated with immune response programs, consistent with the established role of RUNX3 in immune cells. Specifically, RUNX3 has been shown to establish chromatin accessibility at cis-regulatory elements (CREs), which is required for granzyme production and cytotoxic T-cell development [3, 44], supporting the association of Runt-domain binding potentials with immune-related gene programs. To confirm the robustness of the functional associations independently of clustering, gene set enrichment analysis (GSEA) was performed as a complementary approach (Supplement Table A4). Results were largely consistent with those derived from the enriched promoter regions of individual DBD-classes, with co-enriched classes frequently sharing overlapping functional terms. A notable exception was DBD-class 2.2 (Supplementary Figure A5a), which was predominantly associated with immunological pathways in the GSEA, in contrast to the neuronal and membrane-related terms observed in the promoter enrichment analysis. However, these enrichments should be interpreted as suggestive regulatory signatures rather than direct evidence TFs occupancy, given that our analysis is based on predicted binding potentials rather than experimentally determined DNA binding.

To the best of our knowledge, this represents the first systematic genome-wide annotation of all human ENCODE-defined promoters by DBD-class binding potential, demonstrating that the accumulation of DBD-class binding sites within promoter regions, reflecting an increased collective binding potential, can be leveraged to derive functionally interpretable regulatory information, as evidenced by the functional coherence observed across the enriched promoter landscape of the human genome. The observation that DBD-class binding potential profiles cluster promoters into biologically coherent functional groups, as exemplified by the neuronal and membrane-related enrichment of TEAD/TEF-1 binding potentials and the immune response enrichment of Runt-domain binding potentials, establishes TFClassPredict as a broadly applicable tool for the functional annotation of regulatory elements from sequence alone, without a direct need for TF expression or experimental binding data.

### Accessible regulatory regions reveal DBD-class binding potential profiles that recapitulate immune cell differentiation

The DBD-class-associated regulatory regions presented in the previous chapter were predominantly linked to cell-fate decisions covering immune cell development, a finding further supported by the GSEA results, in which immunological development pathways were prominently enriched (Supplementary Figure A5a-d). This prompted us to further investigate DBD-class-specific regulatory differences across immune cell lineages with the aim of reconstructing immune cell differentiation processes. To this end, TFClassPredict was applied to the ATAC-seq dataset of Calderon et al. [45], which profiles chromatin accessibility across 25 primary human immune cell types, with a particular focus on lymphoid cells.

As B-cells, T-cells, and NK-cells share a common lymphoid progenitor, we focused our analysis on naive and immature lymphoid cell types. ATAC-seq profiling of these populations revealed substantial inter-lineage differences in chromatin accessibility (Figure 6a; *N*_B-cell samples_ = 4, *N*NK-cell samples = 5, *N*T-cell samples = 8), suggesting divergent regulatory programs across lineages. To that end, we computed TFBPDs across all open chromatin regions and summarized them for each cell type across ENCODE-defined promoter, proximal enhancer, and distal enhancer regions that showed significantly higher accessibility relative to the other cell types (log_2_FC > 0, *p*_adj_ < 0.05) (Supplement Figure 6a-c, Supplement Table A5; *N*_B-cell regions_ = 36, 356, *N*NK-cell regions = 12, 642, *N*T-cell regions = 19, 918). Pairwise differences in TFBPDs were subsequently calculated to reflect relative TFBP enrichment between lineages, thereby identifying DBD-classes with significant enrichment in regions that are more accessible in each naive population than in the other two.

**Fig. 6.**
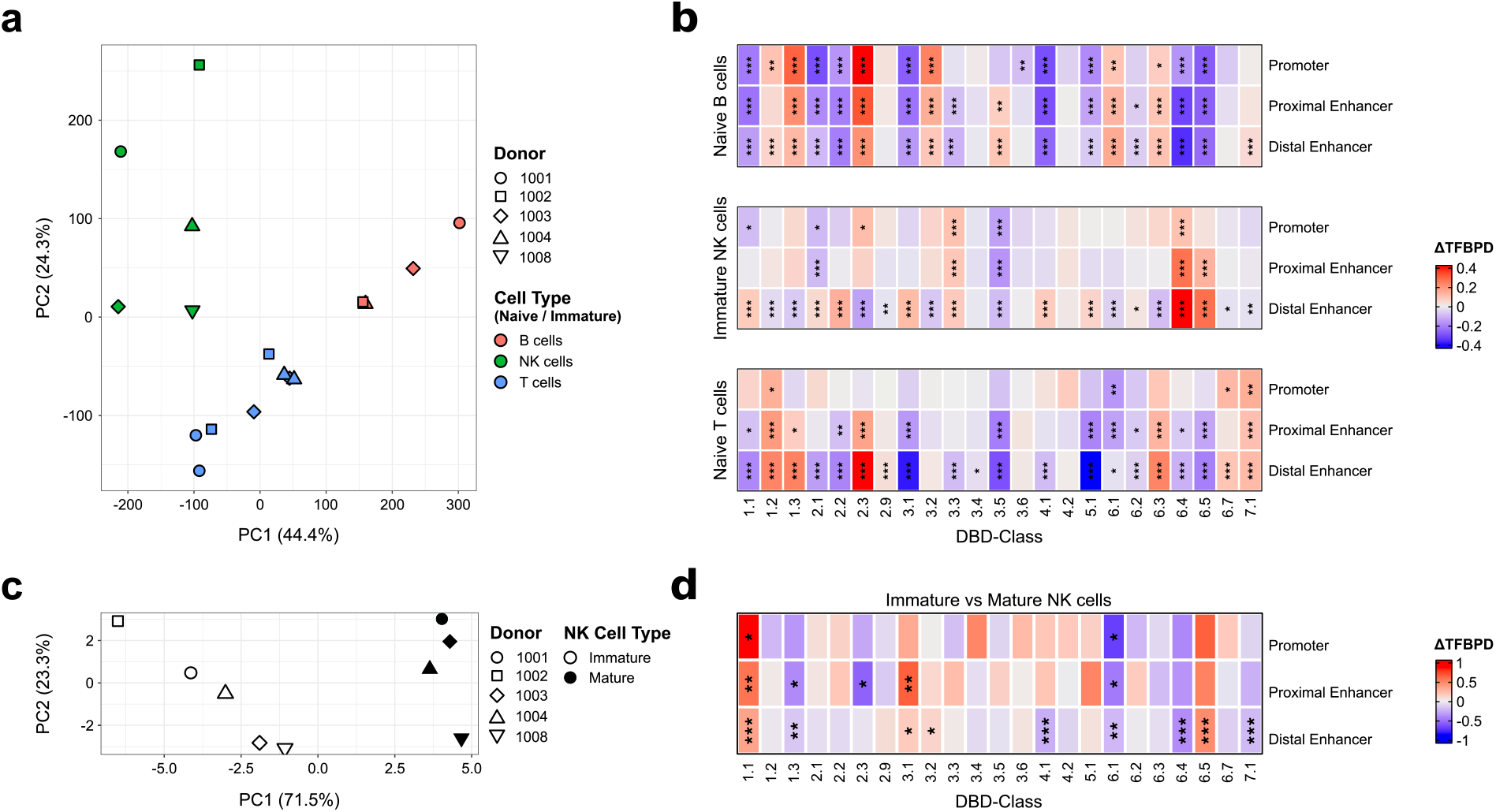
ATAC-seq data from differentiated immune cells reflect well-known biological differentiation processes. **a**: Principal component analysis (PCA) of differentially accessible chromatin regions in naive and immature differentiated lymphoid cells. **b**: Heatmaps of differences in transcription factor binding potential deviations (TFBPDs) derived from differentially accessible regions based on one-versus-other group comparisons. **c**: PCA of differentially accessible chromatin regions in mature versus immature natural killer (NK) cells. **d**: Heatmaps of differential TFBPDs derived from differentially accessible regions in mature versus immature NK cells. Asterisks in heatmaps denote statistical significance of TFBPD en-/derichment: * *p*_*adj*_ < 0.05, ** *p*_*adj*_ < 0.01, *** *p*_*adj*_ < 0.001 based on two-sided t-tests and adjusted within each cell type and set of regions using the Benjamini-Hochberg procedure.

Interestingly, the analysis revealed TFBP enrichment patterns consistent with known immune cell differentiation programs. Particularly, naive B-cell identity and differentiation are known to depend on key TFs, including E2A and TFEB (DBD-class 1.2), EBF1 (DBD-class 6.1), and PAX5 (DBD-class 3.5), all of which direct B-cell fate decisions [4, 46].

On the other side, immature human NK-cell development predominantly depends on TFs such as EOMES and T-bet (DBD-class 6.5), RUNX3 (DBD-class 6.4), ID2 and E2A (DBD-class 1.2), ETS1 (DBD-class 3.5), and E4BP4 (DBD-class 1.1) [2]. Consistent with this, strong TFBP enrichment was observed for Runt factors (DBD-class 6.4) and T-Box factors (DBD-class 6.5), while bZIP factors (DBD-class 1.1) showed preferential enrichment in distal enhancers. The weak enrichment of bHLH factors (DBD-class 1.2) likely reflects ID2-mediated repression of E2A, and the derichment of Trp factors (DBD-class 3.5) is presumably confounded by predominant B-cell binding potentials.

Differentiation into naive CD4^+^ and CD8^+^ T-cells, here combined, is known to be regulated by key TFs, including GATA3 (DBD-class 2.2), EOMES and T-bet (DBD-class 6.5), and Notch (DBD-class 7.1) [47]. While SMAD factor (DBD-class 7.1) enrichment was consistent with the data, the binding potential of DBD-classes 2.2 and 6.5 appeared to be confounded by their predominant activity in NK-cells, where these TFs also play a critical regulatory role [2, 3].

Next, we explored downstream processes related to NK-cell maturation. ATAC-seq data revealed substantial chromatin accessibility changes between immature and mature NK-cells (Figure 6c; *N*Immature NK-cell samples = 5, *N*Mature NK-cell samples = 4) and attributed differential open chromatin regions to either immature NK-cells (log2FC < 0, *p*_adj_ < 0.05) or mature NK-cells (log2FC > 0, *p*_adj_ < 0.05) (Supplement Figure A6d, Supplement Table A5; *N*Immature NK-cell regions = 502, *N*_Mature NK-cell regions_ = 435). Analysis of TFBPs within the differential open chromatin regions for the individual cell types identified strong enrichment (Figure 6d) in distal enhancers for DBD-classes 6.5 and 1.1, suggesting their involvement in NK-cell maturation. Specifically, increased accessibility in DBD-class 6.5 regions in mature NK-cells is consistent with the known role of EOMES and T-bet in driving NK-cell maturation [48], while enrichment of DBD-class 1.1 likely reflects E4BP4 binding activity, a key regulator of NK-cell development [49].

Taken together, the TFBP analysis across naive immune cell lineages demonstrates that TFClassPredict inferred DBD-class binding potentials systematically capture lineage-defining regulatory signatures at chromatin-accessible regions across human immune cell types. The consistent recovery of DBD-classes associated with known lineage-specifying TFs including TFEB, E2A, EBF1, and PAX5 in B cells; T-bet, EOMES, RUNX3, and GATA3 in NK cells; and GATA3, RUNX3, and Notch effects in T-cells, from DBD-class binding potentials derived purely from DNA sequences establishes TFClassPredict as a powerful and biologically principled tool for the interpretation of immune cell regulatory landscapes from ATAC-seq data. More broadly, as TFClassPredict operates exclusively on DNA sequences at the level of individual ATAC-seq peaks, it is directly applicable to both bulk and single-cell ATAC-seq datasets.

## 3 Discussion

Here, we developed TFClassPredict, a DNABERT-based multi-class classification framework that assigns DNA sequences to 23 available DBD-classes defined by the TFClass hierarchy [22], trained on high-confidence directly bound TFBSs from the UniBind repository [29]. We demonstrate that DBD-class sequence grammar is both biologically coherent and computationally learnable from 21bp binding site sequences, and that TFClassPredict achieves robust classification performance, even for underrepresented DBD-classes. We further show that TFClassPredict enhances the reliability of existing PWM-based binding site prediction pipelines and improves three-dimensional genome interaction prediction using Micro-C data, exemplifying its application to ENCODE-defined regulatory elements in the human genome and demonstrating its utility for interpreting ATAC-seq-derived open chromatin regions in human immune cell lineages. Together, these results establish TFClassPredict as a broadly applicable, biologically principled, and practically valuable tool for regulatory sequence analysis. A central conceptual contribution of this work is the demonstration that the DBD-class level of the TFClass hierarchy represents the natural and principled resolution for sequence-based TFBS classification. This argument is supported by two independent and convergent lines of evidence. First, large-scale, systematic characterization of TFBS has established that DBD amino acid sequence similarity directly predicts DNA-binding specificity [23]. We support this principle, which we demonstrate empirically through the k-mer similarity across DBD-classes in our data set. Second, within-class paralog indistinguishability at the PWM level is not merely a technical limitation but reflects a genuine biological reality: paralogs within a class share highly conserved DNA-binding domains that recognize overlapping or identical core sequences, and their binding sites are therefore expected to be similar at the sequence level. Importantly, this conservation of TFBS sequence grammar extends beyond individual TF families to the DBD-class level, where distinct TF families share a common DBD evolutionary origin and frequently interact through homo- and heterodimerization, collectively shaping the binding site grammar of each DBD class. TFClassPredict demonstrates that this level of abstraction is directly learnable and can be effectively applied to identify TFBSs across the human genome.

Despite these advances, several limitations of the current framework should be acknowledged, each pointing to specific directions for future development. First, classification at the DBD-class level, while biologically principled and statistically robust, does not identify which specific TF within a class might bind at a given site. As demonstrated in Figure 4a, b, this limitation can be partially addressed by applying TFClassPredict as a filtering step to PWM-based predictions from tools such as FIMO, which preserves individual TF resolution while substantially reducing false positives. However, sequence information alone is inherently insufficient to determine the binding of a specific TF at a given genomic location, as the dominant binding potential is ultimately determined by the structural properties of the corresponding DBD. More complete disambiguation may be achieved through targeted chromatin profiling experiments such as ChIP-seq or CUT&TAG, or through integration with matched gene and protein expression data, enabling the identification of which specific TF is most likely to occupy a given binding site in a given cellular context. Second, the 21bp sequence window captures the local binding site grammar of each DBD-class, but does not model longer-range sequence context that may influence binding, such as nucleosome positioning, DNA shape, or composite regulatory element environments. This reflects a deliberate design choice rather than a fundamental architectural constraint. Third, while UniBind’s motif-based curation effectively removes indirect binding contamination from the training data, the classifier may underperform on TFs that predominantly engage regulatory elements via tethered or indirect mechanisms. Finally, the current framework covers 23 DBD-classes; as TFClass is explicitly designed to be expandable, future updates incorporating updated DBD-class annotations and additional UniBind training data will further extend the coverage and utility of TFClassPredict.

The core insight of TFClassPredict is that the co-evolutionary relationship between DBD folds and their cognate DNA-binding sequences can be learned from sequence alone. By grouping UniBind TFBSs according to the DBD-based classification of TFClass and training a classifier in DNABERT’s pre-trained k-mer embedding space, TFClassPredict captures the coevolutionary grammar of protein-DNA recognition at the DBD-class level. This conceptually novel and broadly applicable framework achieves solely classification performance, biological interpretability, and strong practical utility that existing PWM-based and deep learning approaches cannot match.

## 4 Methods

### 4.1 Transcription factor binding site sequence data

The UniBind database [29], retrieved from the Unibind zenodo repository (2025 release) [50], was used as the source of TFBS annotations. We specifically used the Direct and Merged Observations (DAMO) collection, which comprises TFBS annotations for 268 human TFs supported by direct experimental evidence. In UniBind, high-confidence TFBSs in the DAMO collection were identified with ChIP-eat analysis pipeline, an entropy-based approach that delineates direct TF–DNA interaction events within ChIP-seq peaks. The use of directly supported TFBSs was intended to minimize noise arising from indirectly bound or spuriously assigned sites. Intersection of the UniBind TF set with TFClass retained 265 TFs with available class annotations.

TFBSs were annotated by the DBD-class of the corresponding bound TF. Genomic region lengths were fixed at 21 bp, in line with the largest binding site defined in the UniBind dataset, to ensure full sequence coverage (Supplementary Figure A1b). Shorter TFBS regions were centered and extended to fit the 21 bps. Underlying genomic sequences were extracted from the USCS hg38 reference genome. To reduce ambiguity in class assignment, TFBSs overlapping with any other TFBS were excluded (Supplementary Figure A1c), and TFBSs located on chromosomes X and Y were removed to avoid potential confounding effects related to sex-chromosome-specific sequence composition and representation. After filtering, the final dataset contained 5,567,690 TFBS sequences. The dataset was partitioned into training and test sets at an 80:20 ratio using a chromosome-stratified random split. The training set was subsequently further sub-divided into training and validation sets using the same stratification procedure. This strategy was adopted to preserve chromosomal representation across all data partitions while maintaining a strict separation between model development and final performance evaluation.

### 4.2 TFClass derived DBD-classes

The TFClass database (2018 release) [22] was utilized to obtain structural domain classifications of TFs. TFClass organizes TFs into a five-level hierarchy: superclass, class, family, subfamily, and genus, based on structural similarity. The Class-level categorization was used as DBD-class labels. DBD-class 0.0 was excluded because of its undefined status within the hierarchy. Classes within superclasses 8 and 9 were not considered hence no human TFBSs data were available for these categories in the UniBind database of TF binding profiles. The same is true for the following classes: 2.5, 2.6, 2.7, 2.8, 3.7, 5.3, 6.6, 7.2. Finally, 23 distinct classes were analyzed. A comprehensive overview of those can be found in the Supplementary Table A1.

### 4.3 Motif to DBD-class mapping

Ensembl IDs for motifs included in the HOMER motif database were manually retrieved from the associated outlinks to TF GeneCards (v5.23) entries provided on the HOMER motif database, accessed on 13.01.2025. Motifs lacking a functional GeneCards outlink were excluded. The DBD-class hierarchy was retrieved from the TFClass website in Turtle format. Ensembl IDs were subsequently used to retrieve the corresponding TFClass annotations from the DBD-class hierarchy via a custom Python script that leverages the RDFLib (v7.1.2) [51] package. Motifs corresponding to TFs belonging to multiple TFClass rank groups due to the presence of multiple DBDs were excluded. The final mapping is provided in the Supplementary Table A1.

Non-redundant JASPAR CORE (2026 release) motifs were manually downloaded from the JASPAR database [52]. Motif-to-DBD-class mapping was retrieved by querying the JASPAR database for TFClass annotations using pyjaspar (v4.0.0) [53]. Annotations were assigned to DBD-classes using a manually curated correspondence table to account for typographical inconsistencies in the TFClass annotations provided by JASPAR (Supplementary Table A1). Motifs that did not map uniquely to a single DBD-class, or for which no human DBD-class annotation was available, were discarded.

### 4.4 Evaluation of conventional TFBP prediction tools

HOMER (v5.1) [32] was used to perform motif enrichment analysis on groups of sequences per DBD-class within the previously defined test set, using the `findMotifsGenome.pl` script with `-size` set to `given`, and the USCS hg38 human genome as reference. The background set is automatically defined by HOMER by randomly sampling GC-content and length-matched genomic sequences. Identified motifs were mapped to DBD-classes according to Section 4.3. Motifs corresponding to the same DBD-class as the query sequence set were labeled as correct, while all others were labeled as incorrect. FIMO [33] was applied using the [MEME Suite Docker image](https://hub.docker.com/r/memesuite/memesuite) (v5.5.9) to perform sequence-wise predictions on the test set using the non-redundant JASPAR CORE motif set. A background model was generated as recommended by the FIMO authors using the `fasta-get-markov` function with default parameters. Predictions were filtered by *p* < 0.001 and mapped to DBD-classes as described in Section 4.3. As multiple predictions are retrieved per sequence, a DBD-class was considered called when at least one motif belonging to that class was identified given the applied filtering criteria. Post-hoc filtering using TFClassPredict was performed by querying each sequence independently with both FIMO and TFClassPredict. Sequences for which the two methods yielded diverging DBD-class assignments were subsequently removed.

### 4.5 TFClassPredict architecture evaluation

To establish TFClassPredict, DNABERT [15] and DanQ [12] were systematically evaluated as candidate model architectures. DNABERT is a BERT-based transformer model pretrained on the entire human genome, comprising 12 transformer layers with multi-head self-attention mechanisms, operating on k-mer tokenized sequences, with four model variants available differing in k-mer size (3-6). DanQ is a hybrid deep learning framework combining a CNN for motif detection with a BLSTM layer to capture regulatory grammar governing the spatial arrangement and co-occurrence of sequence motifs, taking one-hot encoded sequences as input, whereby each nucleotide is represented as a binary vector of length four with a value of one at the position corresponding to the present base (A, C, G, T) and zeros elsewhere. To contextualize model performance, both architectures were further bench-marked against three baseline models: a NN operating on k-mer tokenized sequences, a NN operating on one-hot encoded sequences to isolate the effect of input representation, and a random baseline implemented using scikit-learn’s (v1.8.0) [54] `DummyClassifier`, which assigns predictions by randomly sampling class labels according to the class-wise prior probabilities derived from the training set, thereby reflecting the expected random chance performance under class imbalance.

Class predictions were formulated as a multi-class classification task with exclusive class membership for each input sequence. As described earlier, the dataset was filtered to retain unique binding sites, ensuring that each sequence mapped to a single label across the 23 DBD-classes. The zero vector represents the absence of binding preference for any class.

DNABERT was fine-tuned for DBD-class level multi-class classification by extending the base model with a task-specific classification head, while allowing all model weights to be updated during training. The classification head comprises a GlobalAveragePooling1D layer, followed by a Dense layer (128 units, ReLU activation, L2 kernel regularization), Dropout (0.5), a second Dense layer (64 units, ReLU activation, L2 kernel regularization), a second Dropout (0.5), and a final Dense output layer with 23 units corresponding to the number of DBD-classes. Input sequences of 21 bps were split into overlapping k-mers and tokenized using the BERT tokenizer from the HuggingFace transformers (v4.46.3) package [55]. As DNABERT provides separate pretrained models for different k-mer sizes, the k-mer size was treated as a hyperparameter and optimized on the validation set. The best-performing configuration, using 6-mers, was selected for all subsequent analyses (Supplementary Table A2). To establish a challenging baseline, a NN of similar structure was constructed, comprising a Dense layer (5000 units, ReLU activation, L2 kernel regularization), Dropout (0.5), a second Dense layer (2500 units, ReLU activation, L2 kernel regularization), Dropout (0.5), and a final Dense output layer with 23 units, using the same 6-mer tokens as DNABERT as input. The number of units was deliberately chosen to be large, ensuring sufficient representational capacity to process the high-dimensional k-mer input space and establishing a competitive rather than underpowered baseline.

DanQ was reimplemented according to the original publication [12], except for the Conv1D layer kernel size, which was reduced to 5 to accommodate substantially shorter 21 bp input sequences compared to the 1000 bp sequences used in the original study. In comparison to DNABERT, DanQ relies on one-hot-encoded sequences as input. To compare the two input types, an analogous challenging baseline NN was constructed with the same structure as the 6-mer-based baseline, except that one-hot-encoded sequences were used, similar to DanQ, as input in place of 6-mer tokens.

All models employed a softmax activation function in the final layer and were trained using the Adam optimizer with cross-entropy loss for up to 60 epochs at a learning rate of 0.001. Early stopping was applied as a regularization criterion, terminating training when validation loss did not improve for 3 consecutive epochs. All models were implemented and trained using TensorFlow (v2.13) [56].

Finally, per-DBD-class classification thresholds were independently optimized for each class by maximising the class-wise F1 score, and model performance was assessed using the following metrics:

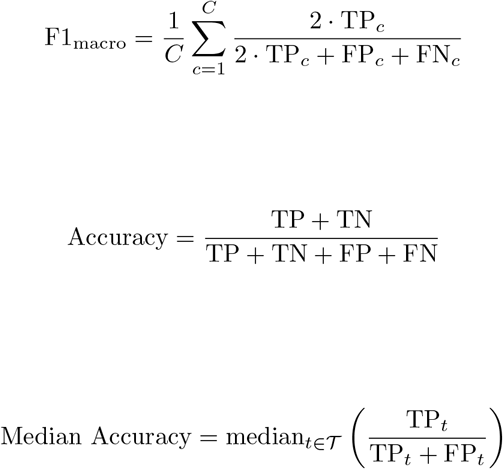

where TP, TN, FP, and FN denote true positives, true negatives, false positives, and false negatives, respectively, and *T* denotes the set of all TFs and *C* describes the number of DBD-classes.

After final model selection, TFClassPredict was extended with a 1 bp sliding window approach to scan sequences larger than the model’s input window, with the resulting TFBPs subsequently summed per region.

### 4.6 Micro-C interaction count prediction

To estimate TFClassPredict’s performance in predicting chromatin interaction counts, Micro-C and ATAC-seq data were retrieved for two cell lines from the 4D Nucleome Data Portal [57, 58]. For HFFc6, files `4DFNIPC7P27B.hic` and `4DNFIWQJFZHS.bb` were used, downloaded on 10.02.2025, and for `H1-hESC`, files `4DNFI2TK7L2F.hic` and `4DNFI247OOFU.bb`, downloaded on 23.07.2025.

TAD were called for each cell line from the respective Micro-C file using the `arrowhead` function from the Juicer platform (v1.6) [59] with default parameters, 5 kbp resolution, and Knight-Ruiz matrix balancing for normalization. ATAC-seq peaks were associated with the genomic bins of the Micro-C matrix, with bins being left-inclusive and right-exclusive. If ATAC-seq peaks cross bin boundaries, they were associated with both bins. TADs with fewer than two ATAC-seq peak-associated bins were disregarded. Subsequently, the Micro-C Knight-Ruiz-balanced interaction counts for all possible combinations of ATAC-seq peak-associated bins within TADs were retrieved from the Micro-C files using the hic-straw (v1.3.1) library [59]. Combinations of bins without an available interaction count value (“NaN”) were discarded. To prevent data leakage, bin combinations were deduplicated. In total, 196,713 bins were obtained for HFFc6 and 189,768 bins for H1-hESC. TFBPs determined by either HOMER or TFClassPredict from sequences pre-selected via ATAC-seq peaks were aggregated by the DBD-class level. TFBP predictions for ATAC-seq peaks were grouped by TAD genomic bins and summed, stratified by the class.

Random Forest (RF) models were trained to predict interaction counts between two bins using the concatenated count vectors as inputs. Model training was performed with five-fold cross-validation using the scikit-learn `RandomForestRegressor`, configured with 500 estimators, bootstrapping enabled, and 200,000 samples drawn per estimator. To account for the power-law relationship between interaction frequency and genomic distance, linear regression models were first fitted to log-transformed distance and interaction count values on the training folds and applied for inference on the validation folds. RF models were subsequently trained to predict the resulting residuals, allowing the assessment of TFBP influence on interaction counts independently of the linear genomic distance. Final interaction count predictions were obtained by summing the predicted residual and the fitted linear regression value, followed by retransformation with a correction term of half the training residual variance to account for Jensen’s inequality.

Model performance was evaluated using the Pearson correlation coefficient for both the predicted residuals and the genomic binned interaction count predictions, with the latter additionally stratified by genomic distance to account for its strong predictive influence.

### 4.7 Transcription factor binding potential deviation

To quantify the enrichment of DBD-class specific binding potentials for single genomic regions, we computed transcription factor binding potential deviation (TFBPD) scores. We first constructed a contingency table **X** ∈ ℕ ^*G*×*D*^, where rows correspond to genomic regions (*G*) and columns to DBD-classes (*D*), with each entry *x*_*gd*_ representing the number of TFBPs identified within genomic region *g* for DBD-class *d*. For each entry, we computed the expected value *e*(*x*_*gd*_) under a null model preserving the marginal distributions of binding potentials per region and per DBD-class:

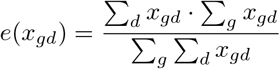

The TFBPD score for each region-DBD-class pair was then computed as the normalized deviation of the observed from the expected value, inspired by ChromVAR’s [60] accessibility deviations:

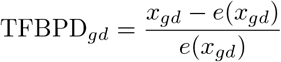

A positive TFBPD score indicates that a genomic region exhibits more TFBPs for a given DBD-class than expected by chance, suggesting specific regulatory enrichment. Since TFBPD scores are normalized by the expected value derived from the marginal distributions, they account for differences in region length and the prior distribution of binding potentials across DBD-classes, correcting for the tendency of certain DBD-classes to be called more frequently. To further enable comparability across DBD-classes and improve numerical properties for downstream analysis, TFBPD scores were subsequently z-scored within each DBD-class.

### 4.8 Genome-wide analysis of TFBPs

To evaluate TFClassPredict genome-wide, we generated genome tracks for the UCSC hg38 reference genome. The human genome was k-merized into unique 21 bp k-mers, stored in a Lightning Memory-Mapped Database (LMDB), and annotated with TFClassPredict predictions. Sequences containing more than a 6-mer of N’s, were discarded. Genomic tracks were subsequently generated for every 21 bp window by scanning the human genome and retrieving the corresponding k-mer annotations from the database. These precompiled predictions, which substantially improve performance for downstream analyses, are publicly available via Zenodo [61].

To investigate the distribution of TFBPs within cis-regulatory elements (CREs), we utilized the ENCODE4 cCREs registry [62], which classifies genomic regions into seven categories based on epigenomic signatures: promoters, proximal enhancers, distal enhancers, H3K4me3-marked open chromatin regions, CTCF-bound accessible sites, open chromatin regions with TFBSs, and TFBS-only regions. We focused our analysis on promoters, proximal enhancers, and distal enhancers, as these regulatory categories offer the highest biological interpretability.

To assess promoters with strong DBD-class-specific enrichment profiles, we computed TFBPDs across the ENCODE4-defined promoter set, retaining only those that ranked in the top 1% for at least one DBD-class. The resulting enriched promoters were embedded using t-SNE and subsequently clustered. To identify the optimal clustering configuration, we evaluated combinations of t-SNE parameters (perplexity ranging from 5 to 55 in steps of 5, and 2 or 3 components) using cosine distance as the similarity metric, alongside two clustering approaches: HDB-SCAN and agglomerative clustering with Ward’s linkage, as implemented in scikit-learn. The configuration yielding the highest silhouette score was selected for final analysis (Supplementary Table A4). Promoters were subsequently linked to their nearest gene using GenomicRanges [63] and TxDb human gene annotations [64], retaining only regions within 50 bp of the gene body. Gene Ontology (GO) overrepresentation analysis per cluster was performed using the `enrichGO` function on gene ontology biological processes from clusterProfiler (v4.16.0) [65]. In addition, Gene Set Enrichment Analysis (GSEA) was performed using the `gseGO` function from the same package. Since several promoters can map to the same gene, we first collapsed scores per DBD-class by retaining, for each gene, the maximum TFBP score across its associated promoters. GSEA was then run against the GO biological process ontology with `nPermSimple = 10,000`. Only gene sets with a significantly positive normalized enrichment score (NES > 0, *p*_*adj*_ < 0.05) were retained for downstream interpretation, reflecting enrichment among genes with the highest TFBP scores.

### 4.9 Immune cell ATAC-seq processing

ATAC-seq data for several immune cell types were obtained from the GEO database (GSE118189), excluding IL-2-treated samples, resulting in a total of 97 samples across 6 donors and 25 cell types. Data were processed with the nfcore/atacseq pipeline (v2.1.2) [66, 67] using the recommended best-practice workflow. Raw paired-end reads were assessed for quality with FastQC and summarized with MultiQC, followed by adapter and quality trimming using Trim Galore. Trimmed reads were aligned to the USCS hg38 human reference genome using BWA-MEM, and alignments were coordinate-sorted with SAMtools. Reads mapping to mitochondrial DNA, low-quality alignments, and duplicate reads were removed as part of the standard pipeline filtering steps. Peaks corresponding to accessible chromatin regions were identified using MACS2. Regions overlapping the ENCODE hg38 blacklist (v3) were excluded from downstream analyses. A consensus peak set was constructed by taking the union of all peaks identified across samples. Unless otherwise stated, default parameters of the nf-core/atacseq pipeline were used.

### 4.10 TFBPs changes in immune cell differentiation

To assess changes in TFBPs across immune cell types, differentially accessible regions were identified from the consensus peak set. Regions located on gonosomes, alternative contigs, or exceeding 3000 bp in length were excluded prior to analysis. Low-count peaks were subsequently removed using the `filterByExpr` function from edgeR (v.4.8.0) [68] with default parameters, retaining only sufficiently covered regions for downstream TFBP analysis. Library sizes were normalized using the TMM strategy, and count data were transformed using the voom method from the limma package (v3.66.0) [69]. A linear model was fitted to identify differentially accessible regions, with cell type and donor included as covariates to account for biological and inter-individual variation.

To assess changes in TFBPs between cell types, TFBPDs were calculated for all filtered open chromatin regions. Differences in TFBPDs between conditions were assessed by comparing sets of regions showing significant differential accessibility using Student’s *t*-tests and adjusted using the Benjamini-Hochberg correction. For the three-way comparison between immature NK-cells, naive B-cells, and naive T-cells, differentially accessible regions were identified using *F* -test (two-sided) statistics on the obtained coefficients with Benjamini-Hochberg-corrected *p*_*adj*_ < 0.05, in a one-versus-rest scheme, contrasting each cell type against the mean of the remaining two. Differentially accessible regions were assigned to a given cell type if the log_2_ fold change relative to the background exceeded 0, indicating greater chromatin accessibility in that cell type than in the others. An analogous analysis was performed for the pairwise comparison between immature and mature NK-cells using *t*-statistics (two-sided). A full list of the differential accessibility analysis for both comparisons can be found in the Zenodo repository [61].

### 4.11 AI usage

Claude Sonnet 4.6 (Anthropic) was used exclusively for language editing and refinement of written passages during the preparation of this manuscript. All scientific content, analyses, and conclusions are solely the work of the authors, who take full responsibility for the accuracy and integrity of this work.

## Code availability

The TFClassPredict software is publicly available at GitLab and can additionally be obtained through PyPI.

## Data availability

All data, models, and precompiled genomic tracks associated with TFClassPredict are publicly available on Zenodo [61]. The original UniBind data can be found in the corresponding Zenodo repository [50].

## Declarations

The authors declare no competing interests.

## Author contributions

Conceptualization: C.I., C.H.T., M.H.; Computational analysis: C.I.; Model development: C.H.T., C.I.; Data collection, processing, and curation: C.I. C.H.T., B.C.H., U.A., I.V.H. Data interpretation: C.I., B.C.H., M.H., I.B., T.B., Ressources: I.B., T.B.; Visualization: C.I., B.C.H., U.A., I.V.H; Writing: C.I., C.H.T., M.H.

## Acknowledgements

This research was supported by the KFO 5002 grant of the Deutsche Forschungsgemeinschaft (DFG). We acknowledge support through the Open Access Publication Funds and transformative agreements of the University of Göttingen. We further acknowledge the support of MEDAS, the Medical Data Science Section of CIDAS, for providing an interdisciplinary research environment and infrastructure. T.B., M.H. and C.I. gratefully acknowledge the support of MEDAS. C.I. gratefully acknowledges the support of the Studienstiftung des Deutschen Volkes. Special thanks to Edgar Wingender for his pioneering contributions to transcription factor classification and for his mentorship in gene regulatory biology. C.H.T. and I.V.H acknowledges the support of the Ph.D. program “Genome Science” –International Max Planck Research School. Lastly, we would like to thank the 4DN consortium for making their data publicly available.

