## Supplementary Figures for "DNA-binding domain-aware classification enables systematic annotation of the regulatory genome"

### Appendix A Supplementary Figures

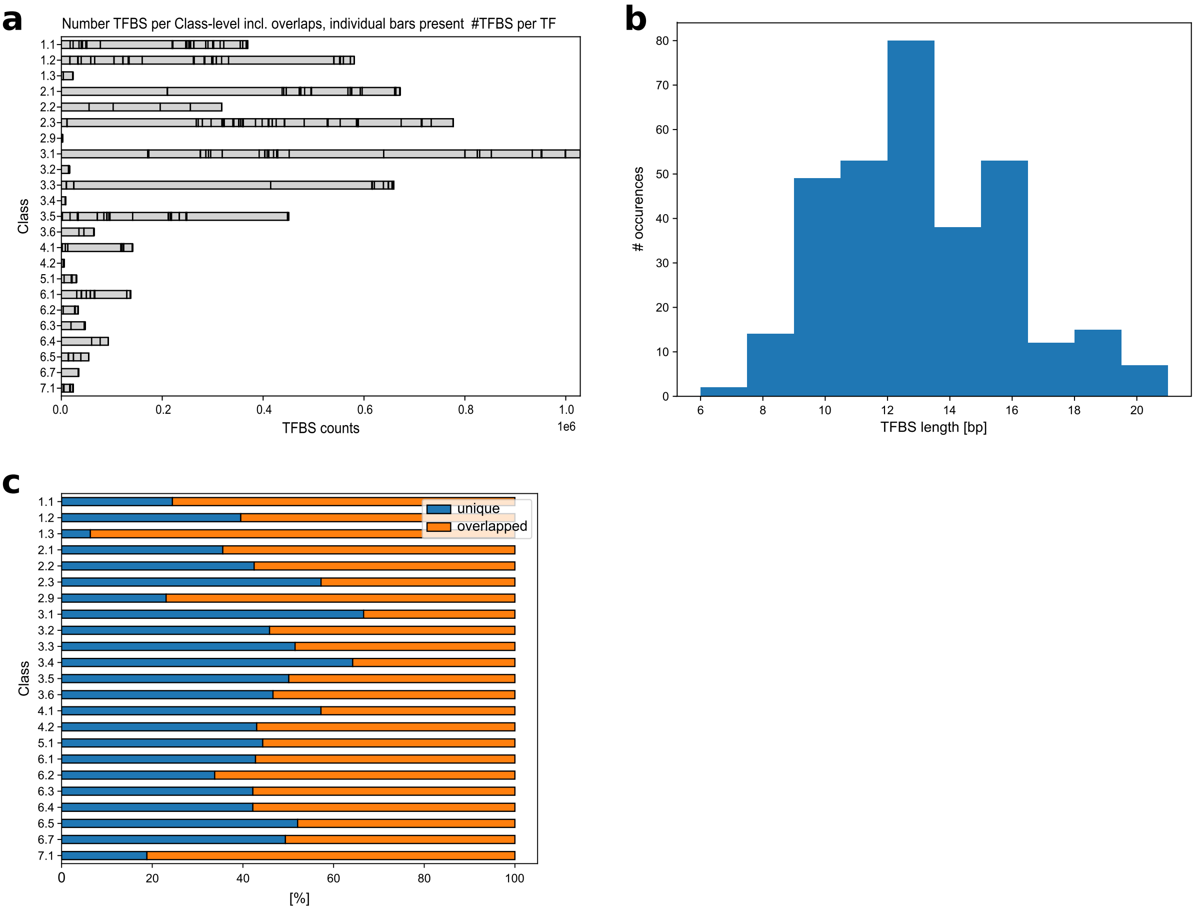


**Fig. A1 Valid TFBSs derived from UniBind. a:** Distribution of TFBSs across DBD-classes in the full UniBind dataset, including overlapping binding sites. Each bar segment represents the number of binding sites for a given TF. **b:** TFBS length distribution across the UniBind dataset. **c:** Percentage of overlapping binding sites removed per DBD-class during construction of the final evaluation dataset.

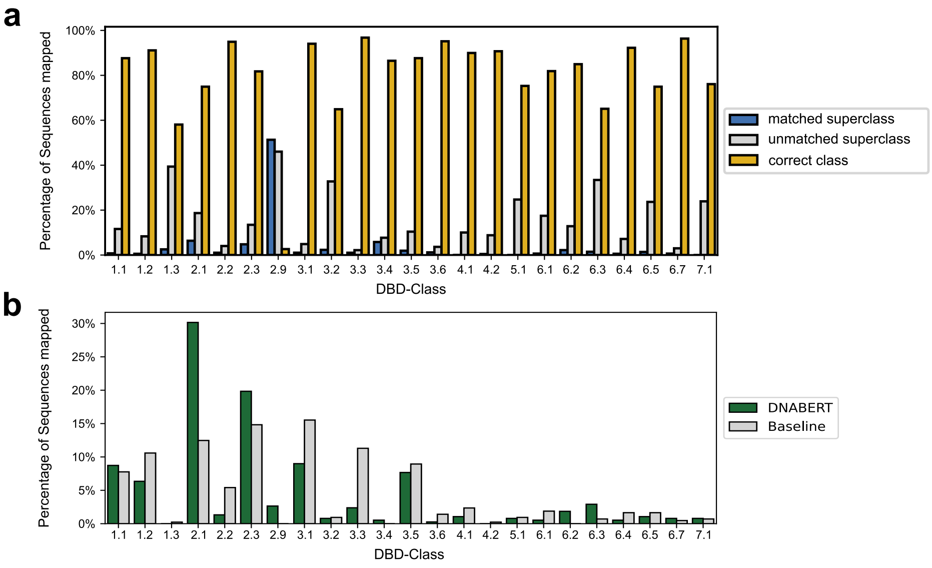


**Fig. A2 DNABERT sequence misclassification results a:** Overview of misclassification rates across all DBD-classes, illustrating the proportion of sequences correctly assigned to their exact DBD-class (yellow) or, where misclassified at the class level, correctly attributed to the appropriate DBD-superclass (blue). **b:** Detailed misclassification analysis of DBD-class 2.9, benchmarked against the class-prior-informed random baseline.

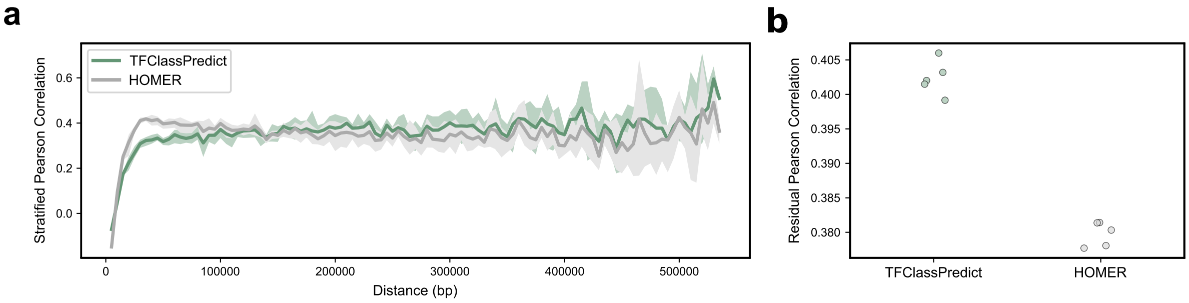


**Fig. A3 Predicting three-dimensional genome interactions from Micro-C data of HFFc6 cells using TFClassPredict-derived transcription factor binding potentials (TFBPs) within ATAC-seq peaks. a:** Pearson correlation between observed and predicted distance-corrected interaction counts for Random Forest (RF) models trained on either HOMER- or TFClassPredict-derived TFBPs. **b:** Pearson correlation stratified by linear genomic distance, comparing RF models based on HOMER- versus TFClassPredict-derived TFBPs.

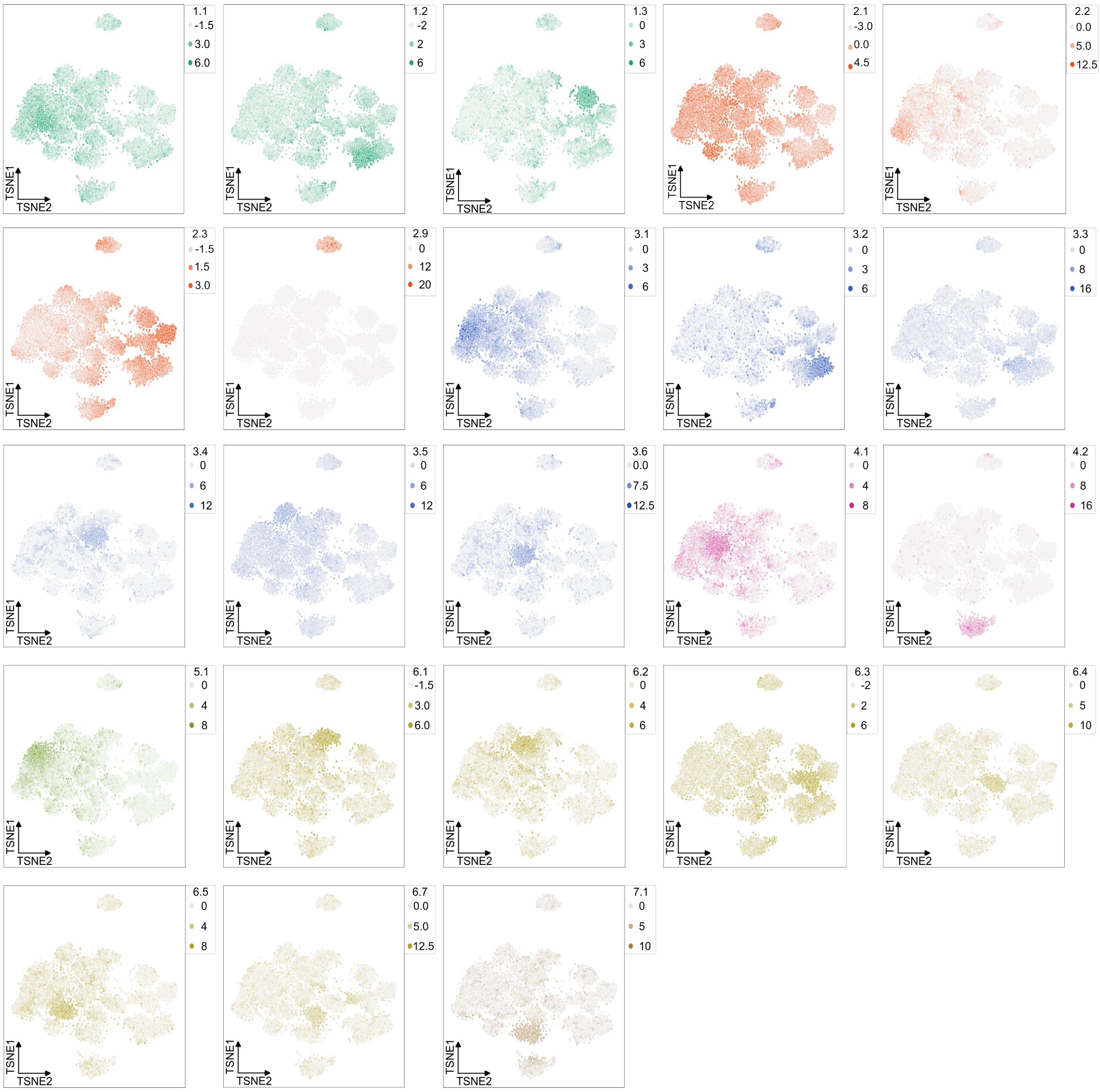


**Fig. A4 t-SNE plot of ENCODE4 cCRE-defined promoter regions with significant transcription factor binding potential deviations (TFBPDs) enrichment in at least one DBD-class.** Each panel shows the distribution of transcription factor binding potential deviations (TFBPDs) for a single DBD-class across the enriched promoter regions.

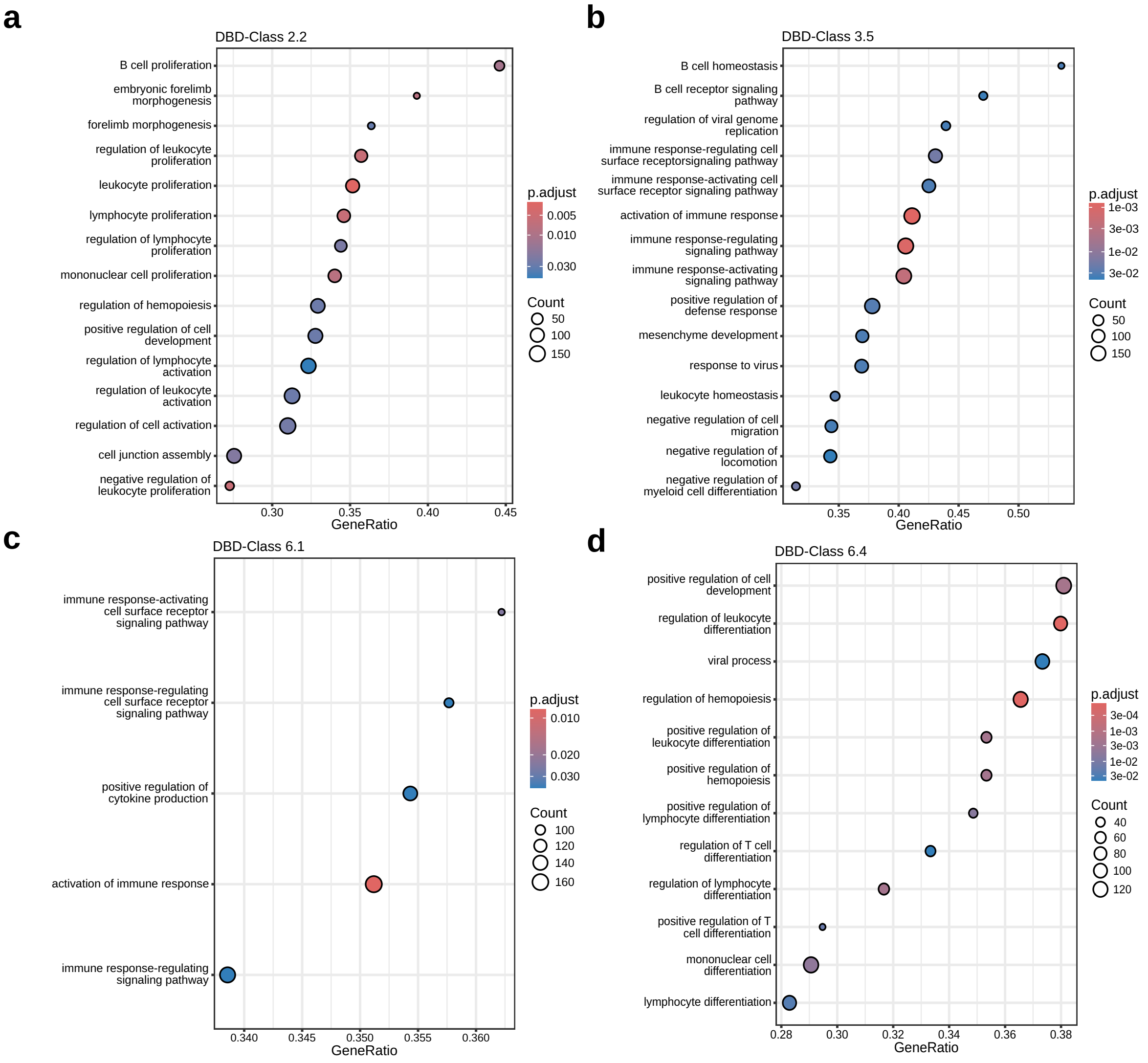


**Fig. A5 GSEA results by DBD-class across all ENCODE4 cCRE-defined promoter regions.** GSEA was performed on GO biological process gene sets, with genes ranked by TFBPDs. Only significantly positive enriched pathways are shown (normalized enrichment score > 0, $p_{adj} < 0.05$, Benjamini-Hochberg correction applied per DBD-class). Plots are restricted to DBD-classes exhibiting associations with immunological pathways: **a:** DBD-class 2.2, **b:** DBD-class 3.5, **c:** DBD-class 5.1, **d:** DBD-class 6.4. The full results can be found in Supplement Table A5.

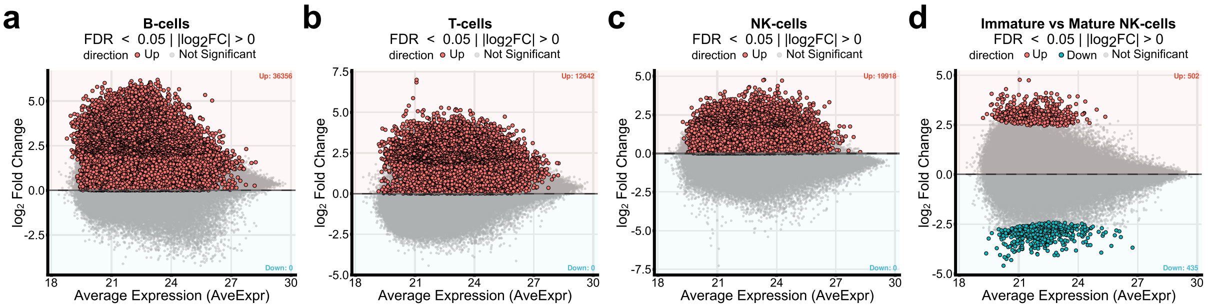


**Fig. A6 MA-plots of differentiated immune cell ATAC-seq data highlighting open chromatin regions used for differential transcription factor binding potential (TFBP) analysis. a:** Regions enriched in naive B-cells (red) relative to immature NK- and naive T-cells. **b:** Regions enriched in immature NK-cells (red) relative to naive B- and T-cells. **c:** Regions enriched in naive T-cells (red) relative to immature NK- and naive B-cells. **d:** Regions enriched in immature NK-cells (blue) and mature NK-cells (red).
